# In-phase and anti-phase dual-site beta tACS differentially influence functional connectivity and motor inhibition

**DOI:** 10.1101/2025.04.28.651005

**Authors:** Tingting Zhu, Alexander T. Sack, Inge Leunissen

**Author notes:** Corresponding author: Email address &.

## Abstract

Inhibitory control relies on coordinated beta-band activity within a fronto-basal ganglia network, which implements inhibition via downstream effects on (pre)motor areas. In this study, we employed dual-site transcranial alternating current stimulation (tACS) targeting the right inferior frontal gyrus (rIFG) and primary motor cortex (M1) to directly manipulate phase relationships in the beta band and assess their effects on both functional connectivity and motor inhibition. Fifty-two healthy participants received in-phase, anti-phase and sham stimulation while performing a stop-signal task. The results revealed that connectivity between rIFG and lM1 increased following in-phase but decreased after anti-phase stimulation. Although no direct modulation of task performance was observed, the greater connectivity increase between the targets during in-phase stimulation was predictive of faster inhibitory performance. In contrast, greater connectivity decreases during anti-phase stimulation were related to faster go responses, suggesting a shift towards less inhibition on the motor system. These findings provide evidence that dual-site beta-tACS can both enhance and impair inhibitory control depending on phase alignment, highlighting its potential as a non-invasive intervention for disorders marked by impaired inhibition.

## Introduction

Motor inhibition is essential for suppressing inappropriate or prepotent responses (Duque et al., 2017; Verbruggen et al., 2019). This process depends on complex neurotransmission within a distributed network of cortical and subcortical regions. Key areas include the right inferior frontal gyrus (rIFG), pre-supplementary motor area (preSMA), and subthalamic nucleus (STN) (Aron, 2007; Aron et al., 2004, 2014; Jahanshahi et al., 2015). Among these, the rIFG is proposed to be the primary initiator of the inhibitory cascade (Duann et al., 2009; Schaum et al., 2020). Together with the preSMA, the rIFG engages the STN via the hyperdirect pathway to initiate motor inhibition. This inhibitory signal is subsequently transmitted to the premotor cortex and the primary motor cortex (M1), where it interacts with ongoing motor preparation to cancel or modify movement execution (Mattia et al., 2012; Mirabella et al., 2011; Stinear et al., 2009). Efficient motor inhibition, therefore, depends on effective long-range communication across this cortico-basal ganglia network.

The communication through coherence hypothesis posits that effective neural communication between brain regions occurs when oscillatory activity of neuronal populations is synchronized in phase (Fries, 2005, 2015). In the context of motor inhibition, increasing evidence suggests that synchronized beta-band activity (15-30 Hz) promotes inhibition. When motor inhibition is experimentally induced using the stop-signal task (SST), the inhibitory network consistently exhibits enhanced beta oscillations (Alegre et al., 2013; Barone & Rossiter, 2021; Castiglione et al., 2019; Swann et al., 2009; Swann et al., 2012; Wagner et al., 2018; Wessel & Aron, 2013; Wessel et al., 2016; Wessel et al., 2019). Transcranial alternating current stimulation (tACS), a non-invasive brain stimulation technique that delivers low-intensity electrical currents to the scalp, is particularly suited for modulating brain oscillations due to its ability to entrain endogenous neuronal activity (Helfrich et al., 2014; Herrmann et al., 2016). Notably, applying tACS at beta frequencies over the M1 has been shown to enhance motor inhibition, establishing a causal relationship between beta activity and inhibitory control (Joundi et al., 2012). However, stimulating M1 in isolation primarily modulates local beta oscillations, providing limited insight into whether beta activity serves as a mechanism for long-range synchronization across the broader motor inhibitory network (Weinrich et al., 2017).

Dual-site tACS (ds-tACS) offers a non-invasive method for modulating inter-regional connectivity by facilitating long-range synchronization between task-related networks (Bramson et al., 2020; Grover et al., 2022; Meijer et al., 2023; Reinhart & Nguyen, 2019). Its effects have been shown to be phase-dependent, such that in-phase tACS (identical currents applied to both regions) enhances functional connectivity, whereas anti-phase tACS (180° phase difference) disrupts it (Helfrich et al., 2014). To date, only a few studies have explored the effects of ds-tACS on motor inhibition, but the findings remain inconclusive (Tan et al., 2022; Fujiyama et al., 2023; Meng et al., 2023).

The current study aimed to systematically investigate whether dual-site in-phase versus anti-phase beta tACS differentially affects functional connectivity in the targeted network to cause or explain individual differences in inhibitory motor control performances. To this end, we applied personalized tACS at the individual beta peak frequency (IBF) using a within-subject design and a high-definition ds-tACS configuration designed to ensure comparable electric field (E-field) distributions across both in-phase and anti-phase conditions. We hypothesized that in-phase beta ds-tACS would enhance functional connectivity within the motor inhibition network and improve inhibitory performance, whereas anti-phase ds-tACS would disrupt these neural dynamics, leading to diminished performance. By implementing a rigorous experimental design, this study aims to advance our understanding of the oscillatory communication mechanisms underlying motor inhibition and to explore the therapeutic potential of ds-tACS as a non-invasive neuromodulator tool for enhancing inhibitory control.

## Methods

### Participants

Fifty-two healthy, right-handed participants (mean laterality quotient: 93.46 ± 1.33% (Oldfield, 1971)), aged 18-36 years (mean age = 22.29 ± 4.21, 18 males) were recruited. Participants met standard eligibility criteria and had no history of neurological or neuropsychiatric disorders or contraindications to transcranial electrical stimulation (Bikson et al., 2009; Woods et al., 2016). All procedures were approved by the Ethics Review Committee Psychology and Neuroscience of Maastricht University, the Netherlands (ERCPN approval code: OZL 204_04_02_2019), and written informed consent was obtained from all participants.

### Stop Signal Task Paradigm

Participants were seated comfortably in a chair approximately 0.6 meters away from the screen (240 Hz refresh rate), with their eyes aligned to the target arrows in the stop-signal paradigm. An anticipated response version of the stop-signal paradigm was used to assess inhibitory control (OSARI, (He et al., 2022)). The task displayed a vertical bar that ascended at a constant speed, reaching the top in 1000 ms (Figure 1A). The target arrows were set at 800 ms after onset. In go trials, the participants were instructed to stop the rising bar at the target arrows as accurately as possible by pressing a button. Go task performance was reinforced by providing feedback. Target arrows changed color to green, yellow, orange or red, indicating the response time within 20, 40, 60 or exceeding 60 ms of the target. In 30% of the trials, the rising bar unexpectedly stopped earlier than the target. These stop trials required participants to withhold their response. The target arrows turned green after successful stops, and red when stopping failed. Stop Signal Delay (SSD) was adjusted to maintain ∼50% stopping accuracy, with staircasing algorithms used for each stimulation condition (in-phase, anti-phase and sham). The initial SSD was set at 600 ms from the trial onset and adjusted in steps of 25 ms. Participants were instructed to aim for green or at least yellow feedback in go trials and to try and withhold the button press on stop trials.

**Figure 1.**
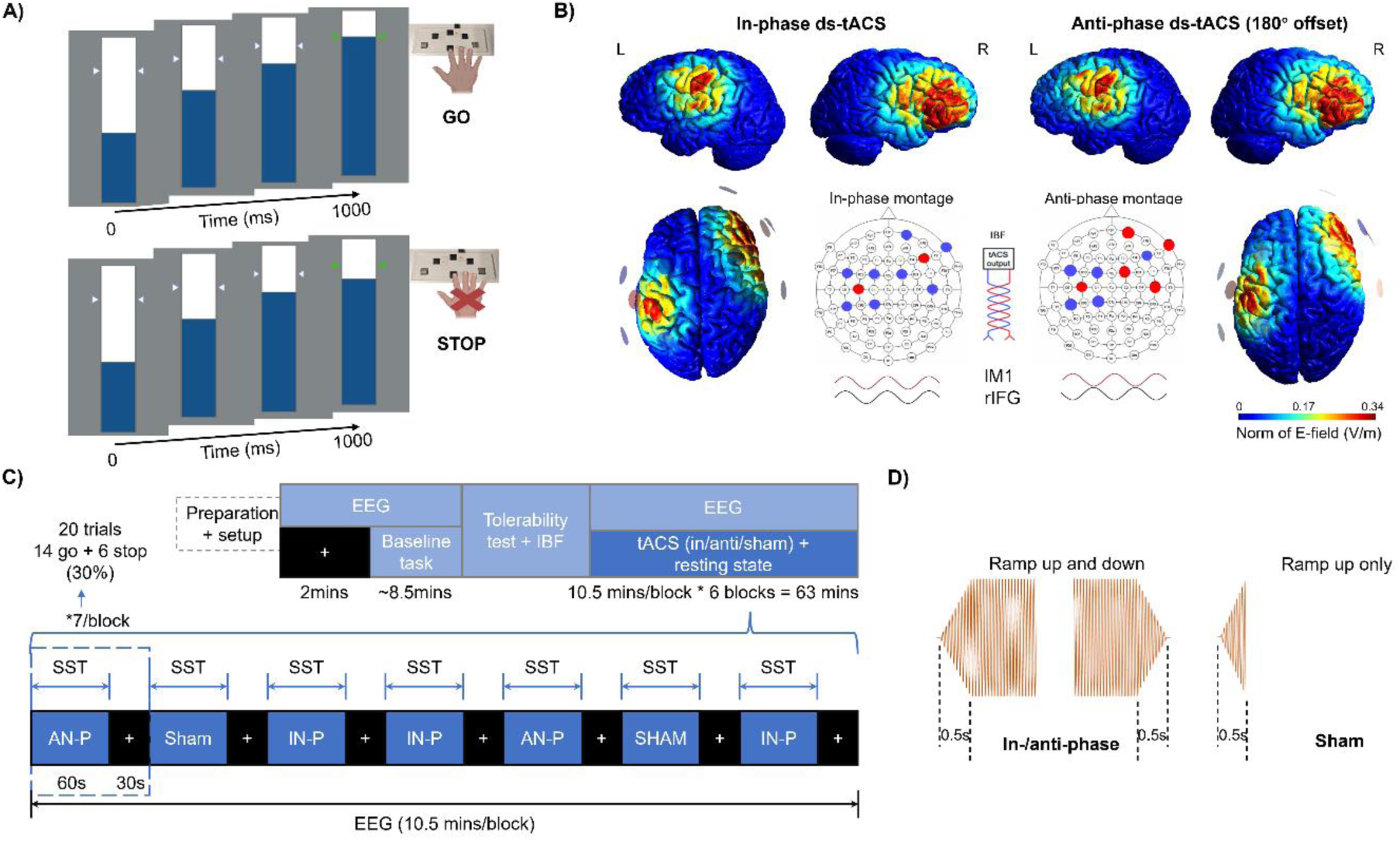
Experimental design for concurrent dual-site tACS (ds-tACS) and EEG study. (**A)** Anticipated response version of stop signal task paradigm. (**B)** The electric field (E-field) modelling, electrode montage and example stimulation waveform for the in-phase (left) and anti-phase (right) conditions. Electrode montage with center electrode (Ø 2 cm) over F6 (rIFG) and C3 (lM1) and surrounding electrodes at ∼5 cm center-to-center distance. Red and blue waveforms illustrate the example phase configurations applied to lM1 and rIFG for each condition. (**C)** Following setup, EEG was recorded during 2 minutes of resting state and ∼8.5 minutes of baseline task without stimulation. Individual beta frequency (IBF) was derived from baseline EEG. Participants then received in-phase (IN-P), anti-phase (AN-P) and sham ds-tACS at their IBF in a counterbalanced order. Each of the six experimental blocks included 7 mini blocks, each with 60 seconds of stimulation and 30 seconds of rest. Each mini block consisted of 20 trials (14 go, 6 stop), preserving a 30% stop trial ratio. **(D)** Example tACS waveform: All stimulation conditions began with a 500-ms ramp-up to ensure gradual onset of current.

Baseline task assessment included 170 trials (51 stops and 119 go trials), with a total duration of about 8.5 minutes. For the task with stimulation, each block contained 140 trials (42 stop and 98 go trials), adding to 840 trials in total (6 blocks, 252 stop and 588 go trials). Correspondingly, 280 trials (84 stop and 196 go trials) were presented per condition. The inter-trial interval ranged from 1.25 to 1.75 seconds.

### tACS-EEG Procedures

Ds-tACS was administered using two NeuroConn stimulators, with waveforms generated via custom Python scripts. The signals were routed via a National Instruments Data Acquisition (NI DAQ) card to the stimulators (Figure 2). Centroid tACS electrodes (2cm Ø) were placed over the rIFG (F6, (Brauer et al., 2018; Cunillera et al., 2016)) and the lM1 (C3). Four surrounding electrodes (2cm Ø) were placed at positions Fp2, FT8, C6 and FC2 for rIFG and FC1, CP1, FC5, and CP5 for lM1 (Figure 1B). The E-field distribution was computed based on standard transcranial direct current stimulation SimNIBS pipeline (www.simnibs.org) (Schwab et al., 2025; Thielscher et al., 2015), applying a 2 mA peak-to-peak current intensity to an example brain (Ernie). All electrodes were modeled as 2 mm-thick rubber layers with a conductivity of 0.1 S/m, and the conductive gel was represented as a 1 mm-thick layer with a conductivity of 3 S/m (as stated by the manufacturer, (Leunissen et al., 2022)). Simulation results indicated that the current flow produced by this 4 × 1 electrode montage was primarily confined to rIFG and lM1 and very similar in the in-phase and anti-phase condition (Figure 1B).

**Figure 2.**
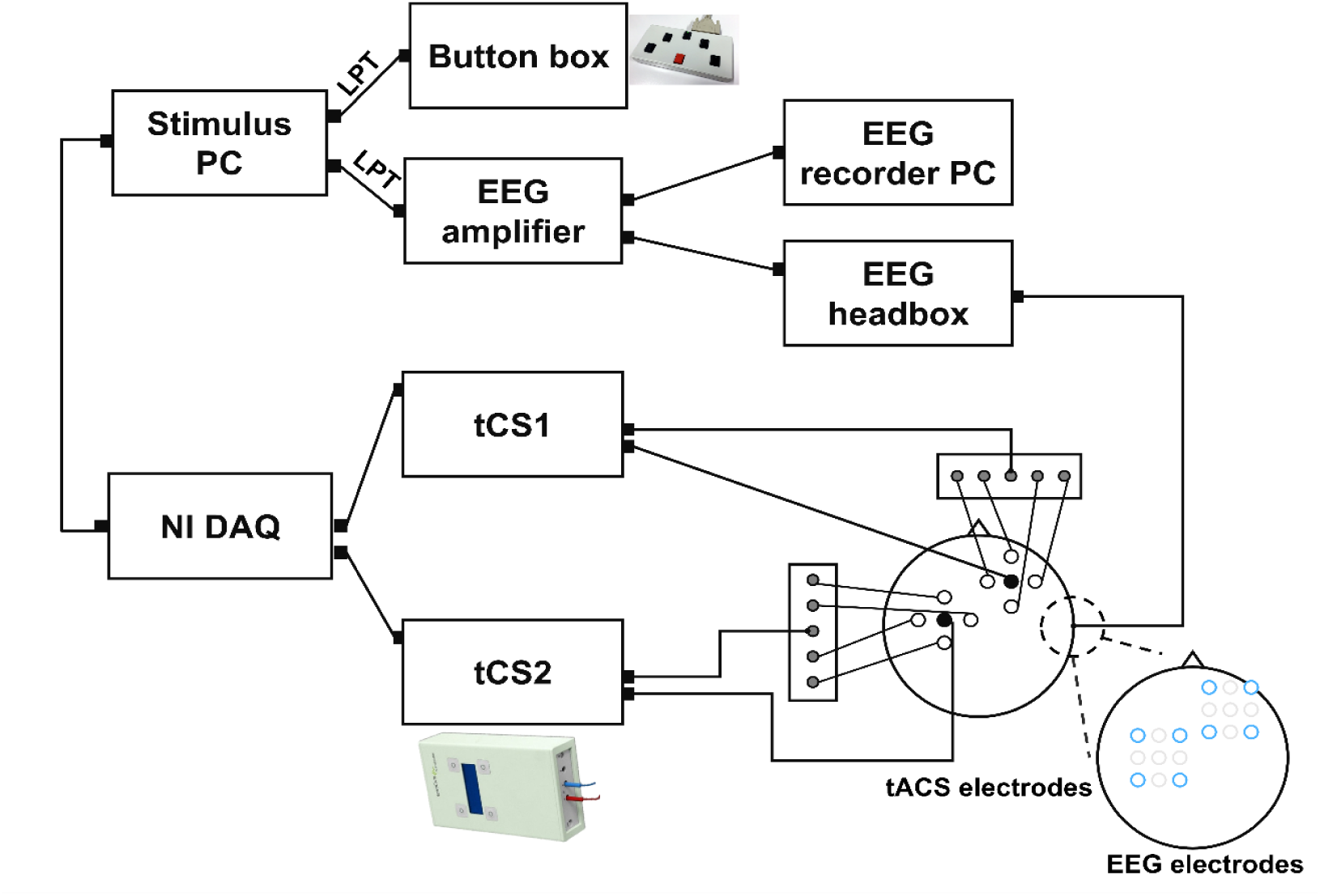
Experimental setup. The tACS waveforms for each stimulation site were generated on the stimulus computer and transmitted to a NI DAQ using custom python scripts, this ensured synchronization between the two stimulation devices. The tACS and EEG electrodes were marked separately for clearer clarification.

EEG was recorded with a BrainAmp DC Amplifier (sampling rate = 5000 Hz, 0.1 mV resolution, 16.4 mV gain). TP10 served as reference electrode, and FPz as ground. EEG signals were recorded from 12 scalp electrodes (TP9, C1, CP3, C5, FC3, F4, AF8, F8, FC6, Fz, FCz and Cz), with additional electrodes placed horizontally and vertically around the eyes to capture eye movement artifacts. Impedance of all EEG and tACS electrodes was kept below 5 kΩ (mean 2.009±0.391 kΩ, range 2.4-5 kΩ).

The experiment began with a 2-minute resting-state EEG (rs-EEG) while participants fixated on a cross, remaining relaxed. This was followed by a baseline SST (∼8.5 minutes), during which IBF was determined (see EEG data analysis). The main experiment involved concurrent ds-tACS and EEG during task performance (Figure 1C). Each participant received three stimulation conditions (in-phase, anti-phase and sham) in a within-subject design. In-phase ds-tACS applied identical currents to lM1 and rIFG, promoting synchronized stimulation; anti-phase involved a 180° phase shift at rIFG to disrupt synchronization. Stimulation intensity was tailored to individual tolerability (mean = 1.917 ± 0.161 mA, range: 1.2–2 mA).

The experiment included 6 blocks (each ∼10.5 minutes), with each block containing 7 mini blocks with 60 seconds of stimulation plus task performance, followed by 30 seconds of rest. Each mini block contained 20 trials (30% stop trials). Stimulation ramped up over 500 ms at onset and ramped down over 500 ms at offset for in-/anti-phase conditions; sham included only the ramp-up (Figure 1D). The stimulation condition order was pseudo-randomized. The onset of stimulation and task were aligned to ensure participants received full-intensity stimulation when the first trial in the mini block started.

Fatigue and stimulation sensations were rated using a 10-point Visual Analogue Scale (VAS) ranging from "not tired/painful at all" to "very tired/painful." Fatigue was also assessed at session start.

### Data Processing

#### EEG Analysis

EEG data from 39 participants were included in the EEG analysis, as EEG recording from the stimulation blocks was not available for 13 participants. EEG processing was performed using FieldTrip (Oostenveld et al., 2011)(v.2017) the following pipeline: 1) removal of bad EEG channels (impedance > 50 kΩ or visually noisy) and interpolation; 2) re-referencing to bilateral TP9/TP10; 3) downsampling from 5000 Hz to 1000 Hz; 4) removal of linear trends and baseline correction; 5) bandpass filtering (1–100 Hz) and removal of 50 Hz line noise (48–52 Hz); 6) regression of eye-related artifacts using the "scrls_regression" function from the EEGLAB AAR plugin (filter order 3; forgetting factor 0.999; sigma 0.01; precision 50) (Gomez-Herrero, 2007); 7) segmentation into 1-second epochs, with visually identified bad trials excluded.

Fast Fourier Transform (FFT) was applied to the data (1-45 Hz frequency range) with a Hanning window and 10-second zero-padding. IBF was determined from successful stop trials in the baseline task using log-transformed EEG spectra from FCz. A linear trend was fitted in a least-squares manner to remove the 1/f property of the spectra (Haegens et al., 2014; Nikulin & Brismar, 2006). The IBF was then obtained by finding the peak from a 3rd order Gaussian curve fitted to the EEG spectra between 15-30Hz (average IBF: 21.09 ± 2.53 Hz, range: 16-26.2 Hz).

EEG data of the rest breaks immediately following stimulation were used to capture the aftereffect of tACS (Figure 1C). For each condition, EEG power was extracted from F8 for rIFG and C1 for lM1. The absolute beta power was averaged within the individual beta frequency window (IBF ± 2 Hz) across trials for each electrode.

To avoid bias in the phase estimation, we selected trials with equal power range for the connectivity analysis (Kovach, 2017). In total, 2.5 ± 1.3 trials with high-power per participant were removed, resulting in 839.1 ± 1.3 trials for anti-phase, 838.3 ± 2.0 trials for in-phase, and 839.3 ± 2.2 trials for sham conditions. After this procedure, there was no significant power difference among ds-tACS conditions anymore (F_(1,38)_ = 0.33, p = 0.722 for lM1; F_(1,38)_ = 1.93, p = 0.150 for rIFG). Then we performed FFT within the frequency range (1 - 45 Hz) using a Hanning window with 2 Hz smooth taps. To comprehensively assess ds-tACS effects on cortical functional connectivity, we applied Phase Locking Value (PLV) and weighted Phase Lag Index (wPLI) within the individual beta frequency window (IBF ± 2 Hz). PLV measures the consistency of phase differences between two signals, reflecting phase synchronization, but is sensitive to volume conduction, potentially inflating connectivity estimates with high type I error (Bruna et al., 2018; Lachaux et al., 1999; Mormann et al., 2005). To mitigate this, wPLI quantifies phase synchronization while excluding zero-lag interactions, offering a more robust estimate of true functional connectivity with potentially high type II error (Vinck et al., 2011). Then the mean connectivity was averaged across trials for each electrode pair.

### Behavioural Analysis

Task performance was estimated based on fifty-two participants. Response times for go trials (goRT) and failed stop trials (fsRT) were determined relative to the trial onset. Early response (> 400ms before the target line) and no response (omission) for go trials were regarded as errors and were removed. The integration method was applied to identify stop signal reaction time (SSRT), in which go omissions were replaced with the maximum RT (1000 ms) (Verbruggen et al., 2019).

### Statistical Analysis

Statistical analyses were conducted using MATLAB 2021b (MathWorks®, Inc., Massachusetts, USA). A two-way repeated measures ANOVA (rANOVA) was used to assess Δpower, with stimulation type (active vs. sham) and region (rIFG vs. lM1) as factors (2x2). Post hoc comparisons were Bonferroni corrected. Paired t-tests were applied to compare ΔPLV, ΔwPLI and ΔRT between the active ds-TACS conditions. One-sample t-tests assessed whether ds-tACS induced significant changes in power, connectivity, and task performance relative to sham. Pearson correlation analyses were performed to examine relationships between Δpower, ΔwPLI, ΔPLV, and Δtask performance. VAS rank values were compared using the Wilcoxon Signed-Rank Test. All tests were considered significant at p < 0.05.

## Results

All participants tolerated the stimulation well and reported only mild itching and warmth underneath the electrode (Table S1).

### Behavioral Performance

A summary of behavioral performance is provided in Table 1. A ∼50% successful stop rate was maintained across all three ds-tACS conditions, with no significant difference observed between them (mean: 49.857 ± 1.598; F_(2,76)_ = 1.205, p = 0.309). As expected, fsRTs were consistently faster than on go trials (goRT) for all participants, in line with the assumptions of the horse-race model (Logan & Cowan, 1984). No significant changes were found in stop-signal reaction time (SSRT) or goRT under either in-phase or anti-phase ds-tACS conditions compared to sham (Figure 3, Table 1).

**Figure 3.**
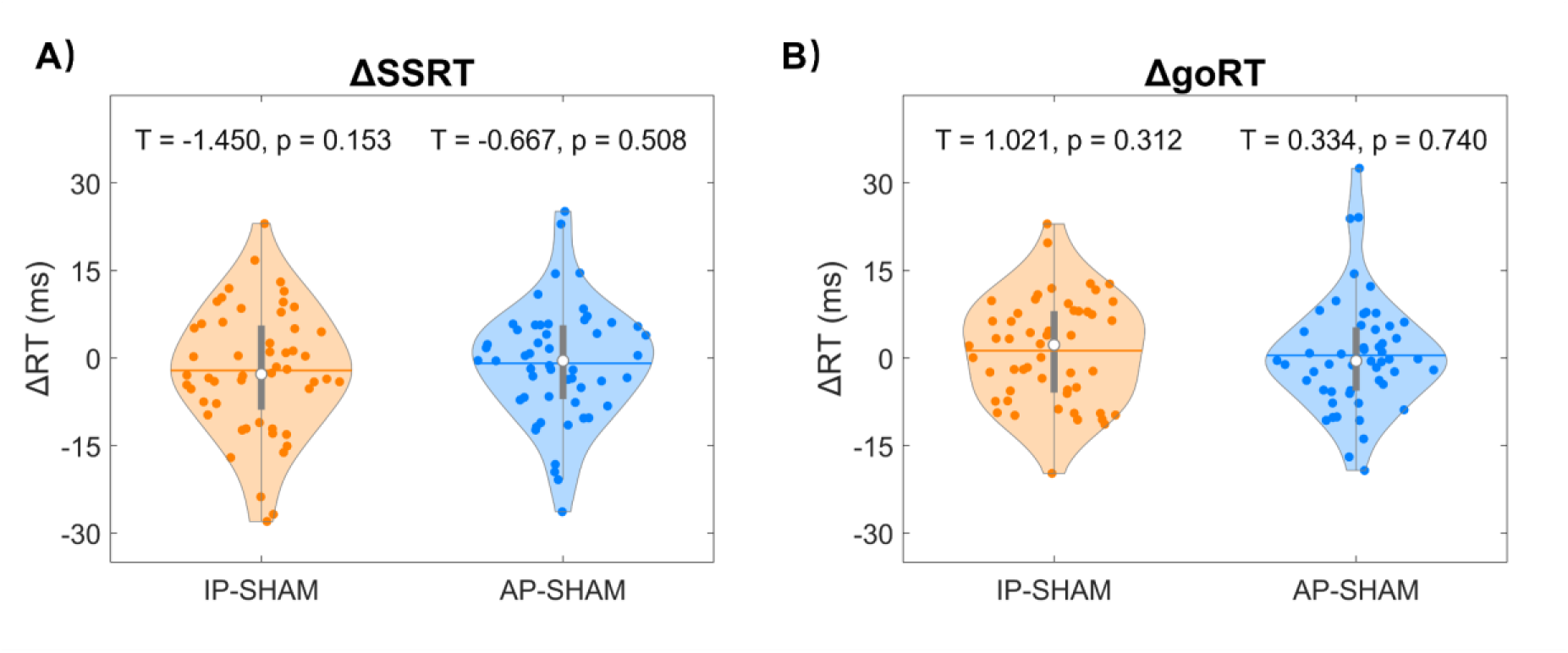
Δ Task performance during ds-tACS for (**A**) stop signal reaction time (SSRT) and (**B**) go reaction time (goRT).

**Table 1.**
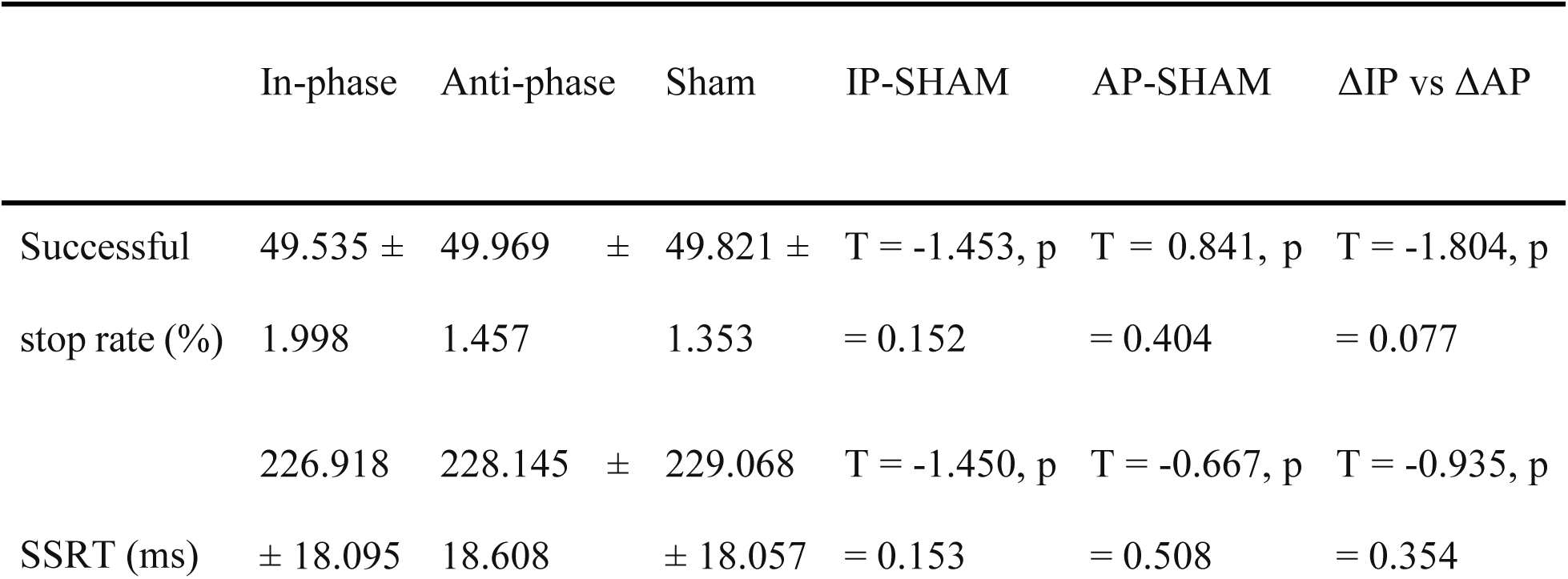
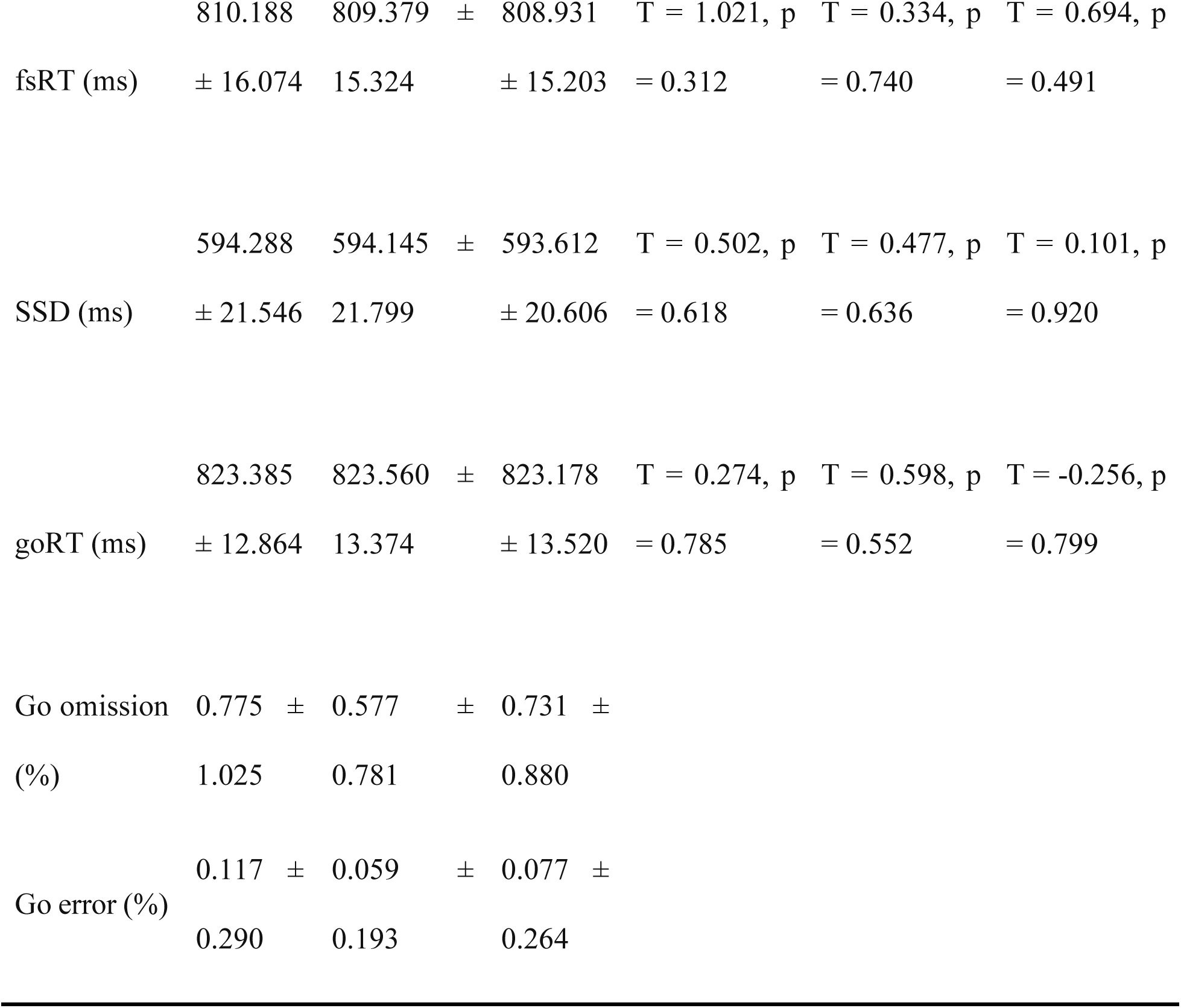
Overview of behavioral performance.

### Beta Power

Both in-phase and anti-phase ds-tACS led to increased beta power in lM1 and rIFG (Figure 4A). A two-way repeated measures ANOVA revealed significant main effects of stimulation type (F_(1,38)_ = 7.267, p = 0.010) and region (F_(1,38)_ = 4.910, p = 0.033), as well as a significant interaction between the two factors (F_(1,38)_ = 6.436, p = 0.015). The stimulation main effect reflected a greater overall beta power increase under in-phase compared to anti-phase tACS (p = 0.010). The regional main effect indicated a stronger power increase in rIFG relative to lM1 (p = 0.033). Post hoc analyses further revealed that in-phase tACS induced significantly greater power in rIFG compared to lM1, and that rIFG power was higher under in-phase than anti-phase stimulation (p = 0.007). Nevertheless, power in both rIFG and lM1 were significantly increased during in- and anti-phase stimulation relative to the sham condition (Figure 4A).

**Figure 4.**
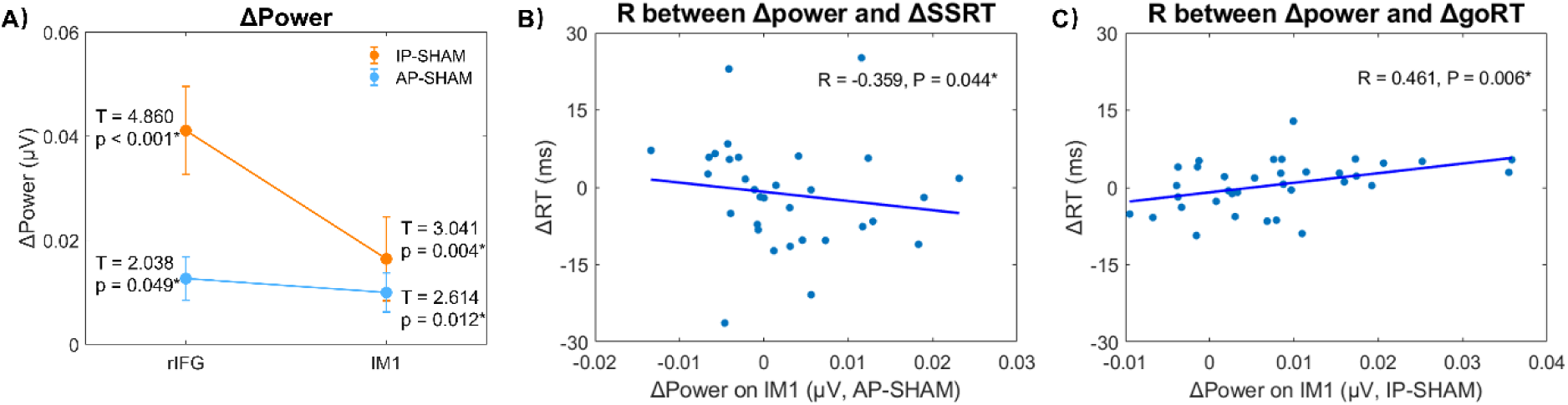
Group results for Δpower. **(A)** ΔPower (mean ± standard deviation). The yellow and blue lines represent the amplitude of Δpower for in- and anti-phase ds-tACS, respectively. T-values and p-values are reported for one-sample t-tests of ds-tACS induced power changes. * p < 0.05. Correlation between Δpower on M1 and **(B)** ΔSSRT in the anti-phase and **(C)** ΔgoRT in the in-phase ds-tACS group.

A greater beta power increase in lM1 under anti-phase ds-tACS was associated with faster motor inhibition performance (R = –0.359, *p* = 0.044), while greater lM1 power increase under in-phase ds-tACS was linked to slower go responses (R = 0.461, *p* = 0.006).

### Phase Locking Value

In-phase ds-tACS significantly increased PLV between rIFG and lM1 (t_(38)_ = 4.356, p < 0.001), whereas anti-phase ds-tACS led to a significant decrease (t_(38)_ = -2.52, p = 0.047), as assessed by one-sample t-tests against zero. A direct comparison revealed a significant difference in PLV changes between the two stimulation types (t_(38)_ = 3.291, p = 0.002; Figure 5A). However, no significant correlation was found between PLV change (ΔPLV) and response time change (ΔRT) induced by tACS.

**Figure 5.**
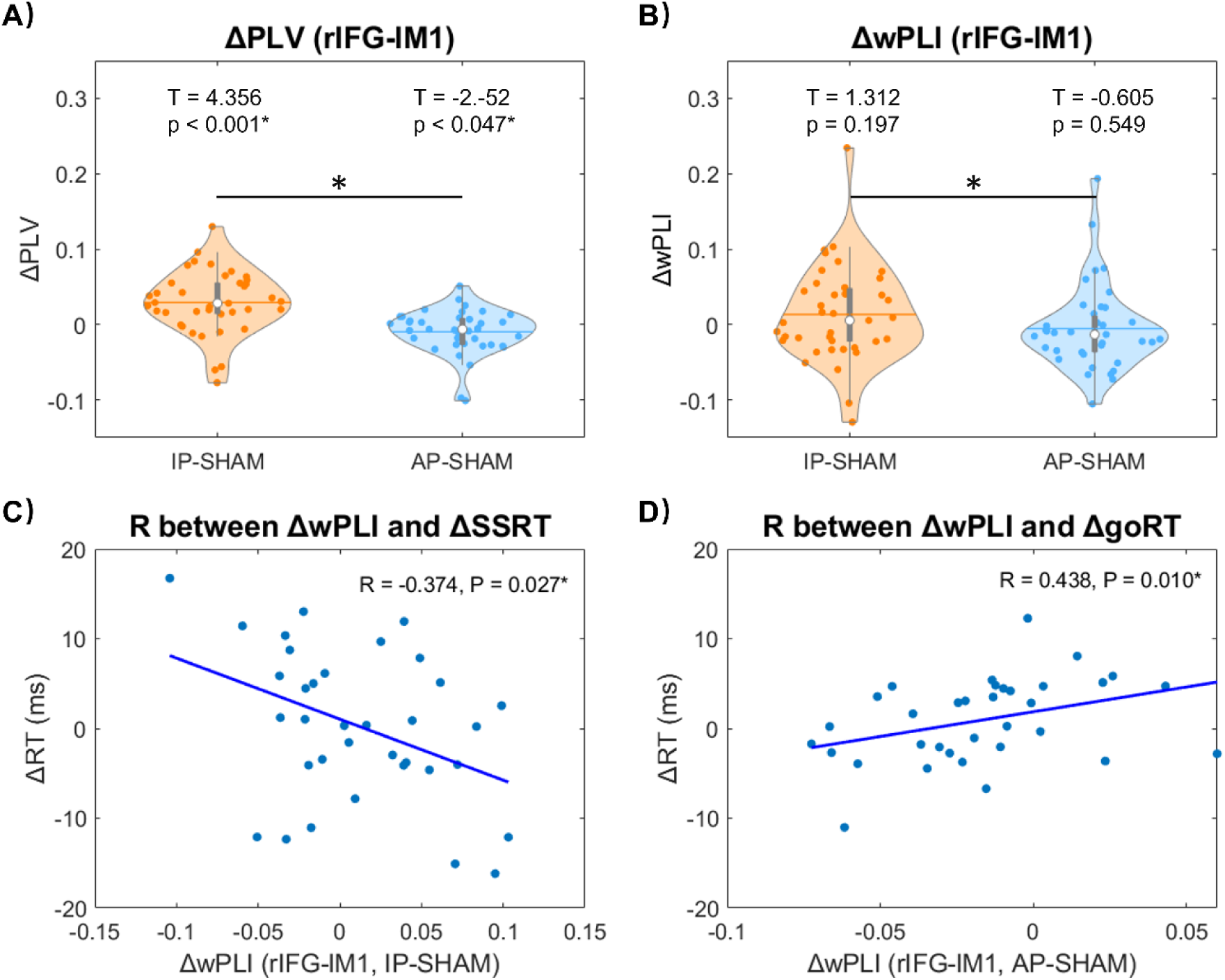
Connectivity changes following ds-tACS. Connectivity comparison across tACS type and connectivity pairs for **(A)** ΔPLV and **(B)** ΔwPLI. T-values and p-values are reported for one-sample t-tests testing ds-tACS induced PLV and wPLI changes. * p < 0.05. The bigger black star in the middle represents a significant connectivity difference between in- and anti-phase tACS conditions. The correlations between ΔwPLI and **(C)** ΔSSRT and **(D)** ΔgoRT.

### Weighted Phase Lag Index

Although ds-tACS did not significantly alter the wPLI between rIFG and lM1 in either condition (in-phase: t_(38)_ = 1.312, p = 0.197; anti-phase: t_(38)_ = -0.605, p = 0.549), a significant difference in modulation between the two stimulation types was observed (t_(38)_ = 2.145, p = 0.038; Figure 5B). Notably, a greater increase in wPLI between rIFG and lM1 under in-phase ds-tACS was associated with faster motor cancellation (R = -0.374, p = 0.027) (Figure 5C). In contrast, under anti-phase ds-tACS, lower wPLI between targets correlated with faster go responses (R = 0.438, p = 0.010) (Figure 5D).

## Discussion

In this study, we employed individualized ds-tACS targeting the rIFG and lM1 to investigate how phase-specific stimulation modulates beta power, interregional connectivity, and motor inhibition. Both in-phase and anti-phase ds-tACS increased beta power at target sites, yet they exerted opposite effects on functional connectivity between rIFG and lM1. In-phase stimulation enhanced phase synchrony and was associated with stronger inhibitory control, while anti-phase stimulation disrupted synchrony and promoted faster motor execution. These findings suggest that manipulating the phase relationship of oscillatory input can selectively influence motor inhibition dynamics by modulating local excitability and long-range connectivity.

### Opposite Modulation of Connectivity and Motor Inhibition by In- and Anti-Phase tACS

In line with previous studies showing that in-phase beta ds-tACS enhances functional connectivity (Fujiyama et al., 2023; Meng et al., 2023), we found that synchronized beta-frequency stimulation increased connectivity between the rIFG and lM1. This enhanced coupling was associated with faster SSRTs, indicating improved inhibitory control. At the same time, in-phase stimulation also enhanced beta power at the stimulation sites, and participants with a greater increase in beta power in M1 displayed slower goRTs, suggesting that heightened beta synchrony imposes a cost on movement initiation. These results align with prior findings that M1 beta activity suppresses motor output (Joundi et al., 2012; Pogosyan et al., 2009) and that stronger functional (Fujiyama et al., 2023; Meng et al., 2023) and structural (Aron et al., 2007; Coxon et al., 2012) connectivity within the motor inhibition network supports more efficient stopping.

In contrast, anti-phase ds-tACS significantly disrupted connectivity between rIFG and lM1, consistent with earlier reports that anti-phase stimulation can interfere with interregional communication (Helfrich et al., 2014). Yet, previous anti-phase ds-tACS studies in the context of motor inhibition did not find a significant decrease in connectivity after anti-phase tACS (Meng et al., 2023; Tan et al., 2022). A key difference in our approach was the use of IBF stimulation, which likely enhanced both the precision and efficacy of neural modulation by aligning with each participant’s intrinsic oscillatory dynamics (Ali et al., 2013; Fang et al., 2024). Additionally, our stimulation was applied online during task performance, in a larger sample and with protocols designed to equate the electric field across in-phase and anti-phase conditions. This design enabled a temporally specific examination of how phase-dependent stimulation influences motor inhibition in the context of ongoing behavior. Crucially, greater reductions in rIFG-lM1 synchrony were associated with faster goRTs, suggesting that decoupling these regions may release inhibitory control over the motor system, thereby facilitating movement initiation.

Taken together, the observed effects on connectivity and behavior seem to reflect a functional trade-off: in-phase ds-tACS reinforces top-down inhibitory signaling at the expense of motor speed, while anti-phase ds-tACS loosens these constraints, promoting quicker and more flexible action control. The finding supports theoretical frameworks proposing that overly synchronized beta activity can rigidify motor control (Brittain & Brown, 2014), while desynchronization may restore adaptability. Future work may explore whether these effects generalize to clinical populations with impaired inhibition, such as individuals with Parkinson’s disease or attention deficit hyperactivity disorder (ADHD), where modulation of beta synchrony could offer therapeutic benefit.

### Enhanced Beta Power after Both In- and Anti-phase ds-tACS

Interestingly, anti-phase stimulation increased local beta power, and within M1 this increase was associated with faster SSRT. This introduces a nuanced picture: while long-range synchrony between rIFG and M1 is necessary for successful inhibition, local increases in beta power may support distinct, perhaps compensatory mechanisms. The interaction between local and network-level dynamics suggests that tACS does not exert uniform effects across spatial scales. Instead, entrainment may strengthen local oscillations while disrupting cross-regional phase alignment, leading to divergent behavioral consequences depending on which mechanism dominates. This complex pattern also helps reconcile seemingly contradictory findings in literature (Seo et al., 2024). For example, prior studies have shown beta power increases during stopping, often interpreted as beneficial (Del Campo-Vera et al., 2022; Leunissen et al., 2022; Schaum et al., 2020; Wagner et al., 2018). Our results suggest that beta power alone is not sufficient—phase alignment across regions is critical for effective top-down control (Helfrich et al., 2014). Disrupting this alignment, even in the context of heightened local beta, can degrade performance. Additionally, the impact of stimulation will also depend on the endogenous phase lag between regions; externally imposed phase relationships can either reinforce or disrupt natural communication dynamics, depending on how closely they match the brain’s intrinsic timing (Elyamany et al., 2025).

### Possible Explanations for Behavioral Effects

There are several plausible explanations for the absence of behavioral improvement. First, motor inhibition is not solely governed by beta synchronization between two regions but likely depends on broader network interactions including the preSMA, STN, and premotor cortex (Aron et al., 2007; Jahanshahi et al., 2015). Our stimulation targeted only rIFG and lM1, possibly omitting critical nodes or pathways necessary for translating oscillatory synchrony into behavioral output. Second, it is possible that beta activity facilitates inhibition only in specific temporal windows or task states. Beta bursts, rather than sustained beta power, have been implicated in the successful cancellation of motor responses (Wessel, 2020). Continuous stimulation at fixed frequencies may not align with these transient, state-dependent dynamics, reducing efficacy. Finally, it is also conceivable that the healthy brain possesses sufficient flexibility or redundancy to compensate for externally induced disruptions in connectivity. Such compensatory mechanisms could buffer behavioral performance, especially when interference affects only a subset of the relevant network, masking the functional impact of targeted modulation.

## Conclusion

Our findings provide compelling evidence that dual-site, phase-specific tACS targeting rIFG and lM1 can differentially modulate beta oscillations and cortical communication in ways that shape motor inhibition. In-phase stimulation enhances beta synchrony and improves inhibitory control, while anti-phase stimulation disrupts synchrony to support faster, more flexible motor execution. These results provide compelling mechanistic evidence that long-range beta-band synchronization plays a critical role in motor inhibition. They also highlight the promise of phase-specific tACS as a targeted therapeutic strategy for disorders characterized by impaired inhibitory control, such as Parkinson’s disease and ADHD.

## Author contribution

T.Z.: Conceptualization, Data Curation, Formal analysis, Investigation, Methodology, Resources, Software, Validation, Visualization, Writing - original draft, Writing - review & editing. A.T.S.: Funding acquisition, Methodology, Supervision, Writing - review & editing. I.L.: Conceptualization, Formal analysis, Funding acquisition, Investigation, Methodology, Resources, Software, Supervision, Validation, Writing - review & editing.

## Data and Code Availability

De-identified raw data and all original code has been deposited at the DataverseNL repository and is publicly available as of the date of publication. Any additional information required to reanalyze the data reported in this paper is available from the corresponding authors upon request.

## References

Alegre, M., Lopez-Azcarate, J., Obeso, I., Wilkinson, L., Rodriguez-Oroz, M. C., Valencia, M., Garcia-Garcia, D., Guridi, J., Artieda, J., Jahanshahi, M., & Obeso, J. A. (2013). The subthalamic nucleus is involved in successful inhibition in the stop-signal task: A local field potential study in Parkinson’s disease. Experimental Neurology, 239, 1–12. 10.1016/j.expneurol.2012.08.027

Ali, M. M., Sellers, K. K., & Frohlich, F. (2013). Transcranial alternating current stimulation modulates large-scale cortical network activity by network resonance. J Neurosci, 33(27), 11262–11275. 10.1523/JNEUROSCI.5867-12.2013

Aron, A. R. (2007). The neural basis of inhibition in cognitive control. Neuroscientist, 13(3), 214–228. 10.1177/1073858407299288

Aron, A. R., Behrens, T. E., Smith, S., Frank, M. J., & Poldrack, R. A. (2007). Triangulating a cognitive control network using diffusion-weighted magnetic resonance imaging (MRI) and functional MRI. J Neurosci, 27(14), 3743–3752. 10.1523/JNEUROSCI.0519-07.2007

Aron, A. R., Robbins, T. W., & Poldrack, R. A. (2004). Inhibition and the right inferior frontal cortex. Trends Cogn Sci, 8(4), 170–177. 10.1016/j.tics.2004.02.010

Aron, A. R., Robbins, T. W., & Poldrack, R. A. (2014). Inhibition and the right inferior frontal cortex: one decade on. Trends Cogn Sci, 18(4), 177–185. 10.1016/j.tics.2013.12.003

Barone, J., & Rossiter, H. E. (2021). Understanding the Role of Sensorimotor Beta Oscillations. Front Syst Neurosci, 15, 655886. 10.3389/fnsys.2021.655886

Bikson, M., Datta, A., & Elwassif, M. (2009). Establishing safety limits for transcranial direct current stimulation. Clin Neurophysiol, 120(6), 1033–1034. 10.1016/j.clinph.2009.03.018

Bramson, B., den Ouden, H. E., Toni, I., & Roelofs, K. (2020). Improving emotional-action control by targeting long-range phase-amplitude neuronal coupling. eLife, 9, e59600. 10.7554/eLife.59600

Brauer, H., Kadish, N. E., Pedersen, A., Siniatchkin, M., & Moliadze, V. (2018). No Modulatory Effects when Stimulating the Right Inferior Frontal Gyrus with Continuous 6 Hz tACS and tRNS on Response Inhibition: A Behavioral Study. Neural Plast, 2018, 3156796. 10.1155/2018/3156796

Brittain, J. S., & Brown, P. (2014). Oscillations and the basal ganglia: motor control and beyond. NeuroImage, 85 Pt 2(Pt 2), 637-647. 10.1016/j.neuroimage.2013.05.084

Bruna, R., Maestu, F., & Pereda, E. (2018). Phase locking value revisited: teaching new tricks to an old dog. J Neural Eng, 15(5), 056011. 10.1088/1741-2552/aacfe4

Castiglione, A., Wagner, J., Anderson, M., & Aron, A. R. (2019). Preventing a Thought from Coming to Mind Elicits Increased Right Frontal Beta Just as Stopping Action Does. Cerebral Cortex, 29(5), 2160–2172. 10.1093/cercor/bhz017

Coxon, J. P., Van Impe, A., Wenderoth, N., & Swinnen, S. P. (2012). Aging and inhibitory control of action: cortico-subthalamic connection strength predicts stopping performance. J Neurosci, 32(24), 8401–8412. 10.1523/JNEUROSCI.6360-11.2012

Cunillera, T., Brignani, D., Cucurell, D., Fuentemilla, L., & Miniussi, C. (2016). The right inferior frontal cortex in response inhibition: A tDCS-ERP co-registration study. NeuroImage, 140, 66–75. 10.1016/j.neuroimage.2015.11.044

Del Campo-Vera, R. M., Tang, A. M., Gogia, A. S., Chen, K. H., Sebastian, R., Gilbert, Z. D., Nune, G., Liu, C. Y., Kellis, S., & Lee, B. (2022). Neuromodulation in Beta-Band Power Between Movement Execution and Inhibition in the Human Hippocampus. Neuromodulation, 25(2), 232–244. 10.1111/ner.13486

Duann, J. R., Ide, J. S., Luo, X., & Li, C. S. (2009). Functional connectivity delineates distinct roles of the inferior frontal cortex and presupplementary motor area in stop signal inhibition. J Neurosci, 29(32), 10171–10179. 10.1523/JNEUROSCI.1300-09.2009

Duque, J., Greenhouse, I., Labruna, L., & Ivry, R. B. (2017). Physiological Markers of Motor Inhibition during Human Behavior. Trends Neurosci, 40(4), 219–236. 10.1016/j.tins.2017.02.006

Elyamany, O., Iffland, J., Bak, J., Classen, C., Nolte, G., Schneider, T. R., Leicht, G., & Mulert, C. (2025). Predictive role of endogenous phase lags between target brain regions in dual-site transcranial alternating current stimulation. Brain Stimul. 10.1016/j.brs.2025.04.011

Fang, Z., Sack, A. T., & Leunissen, I. (2024). The phase of tACS-entrained pre-SMA beta oscillations modulates motor inhibition. NeuroImage, 290, 120572. 10.1016/j.neuroimage.2024.120572

Fries, P. (2005). A mechanism for cognitive dynamics: neuronal communication through neuronal coherence. Trends in Cognitive Sciences, 9(10), 474–480. 10.1016/j.tics.2005.08.011

Fries, P. (2015). Rhythms for Cognition: Communication through Coherence. Neuron, 88(1), 220–235. 10.1016/j.neuron.2015.09.034

Fujiyama, H., Williams, A., Tan, J., Levin, O., & Hinder, M. R. (2023). Comparison of online and offline applications of dual-site transcranial alternating current stimulation (tACS) over the pre-supplementary motor area (preSMA) and right inferior frontal gyrus (rIFG) for improving response inhibition. Neuropsychologia, 191, 108737. 10.1016/j.neuropsychologia.2023.108737

Gomez-Herrero, G. (2007). Automatic artifact removal (AAR) toolbox v1.3 (Release 09.12.2007) for MATLAB. Technology, 3, 1–23.

Grover, S., Wen, W., Viswanathan, V., Gill, C. T., & Reinhart, R. M. G. (2022). Long-lasting, dissociable improvements in working memory and long-term memory in older adults with repetitive neuromodulation. Nat Neurosci, 25(9), 1237–1246. 10.1038/s41593-022-01132-3

Haegens, S., Cousijn, H., Wallis, G., Harrison, P. J., & Nobre, A. C. (2014). Inter- and intra-individual variability in alpha peak frequency. NeuroImage, 92(100), 46–55. 10.1016/j.neuroimage.2014.01.049

He, J. L., Hirst, R. J., Puri, R., Coxon, J., Byblow, W., Hinder, M., Skippen, P., Matzke, D., Heathcote, A., Wadsley, C. G., Silk, T., Hyde, C., Parmar, D., Pedapati, E., Gilbert, D. L., Huddleston, D. A., Mostofsky, S., Leunissen, I., MacDonald, H. J., . . . Puts, N. A. J. (2022). OSARI, an Open-Source Anticipated Response Inhibition Task. Behav Res Methods, 54(3), 1530–1540. 10.3758/s13428-021-01680-9

Helfrich, R. F., Knepper, H., Nolte, G., Struber, D., Rach, S., Herrmann, C. S., Schneider, T. R., & Engel, A. K. (2014). Selective modulation of interhemispheric functional connectivity by HD-tACS shapes perception. PLoS Biol, 12(12), e1002031. 10.1371/journal.pbio.1002031

Herrmann, C. S., Murray, M. M., Ionta, S., Hutt, A., & Lefebvre, J. (2016). Shaping Intrinsic Neural Oscillations with Periodic Stimulation. J Neurosci, 36(19), 5328–5337. 10.1523/JNEUROSCI.0236-16.2016

Jahanshahi, M., Obeso, I., Rothwell, J. C., & Obeso, J. A. (2015). A fronto-striato-subthalamic-pallidal network for goal-directed and habitual inhibition. Nat Rev Neurosci, 16(12), 719–732. 10.1038/nrn4038

Joundi, R. A., Jenkinson, N., Brittain, J. S., Aziz, T. Z., & Brown, P. (2012). Driving oscillatory activity in the human cortex enhances motor performance. Curr Biol, 22(5), 403–407. 10.1016/j.cub.2012.01.024

Kovach, C. K. (2017). A Biased Look at Phase Locking: Brief Critical Review and Proposed Remedy. IEEE Transactions on Signal Processing, 65(17), 4468–4480. 10.1109/Tsp.2017.2711517

Lachaux, J. P., Rodriguez, E., Martinerie, J., & Varela, F. J. (1999). Measuring phase synchrony in brain signals. Hum Brain Mapp, 8(4), 194–208. 10.1002/(sici)1097-0193(1999)8:4<194::aid-hbm4>3.0.co;2-c

Leunissen, I., Van Steenkiste, M., Heise, K. F., Monteiro, T. S., Dunovan, K., Mantini, D., Coxon, J. P., & Swinnen, S. P. (2022). Effects of beta-band and gamma-band rhythmic stimulation on motor inhibition. iScience, 25(5), 104338. 10.1016/j.isci.2022.104338

Logan, G. D., & Cowan, W. B. (1984). On the ability to inhibit thought and action: A theory of an act of control. Psychological Review, 91(3), 295–327.

Mattia, M., Spadacenta, S., Pavone, L., Quarato, P., Esposito, V., Sparano, A., Sebastiano, F., Di Gennaro, G., Morace, R., Cantore, G., & Mirabella, G. (2012). Stop-event-related potentials from intracranial electrodes reveal a key role of premotor and motor cortices in stopping ongoing movements. Front Neuroeng, 5, 12. 10.3389/fneng.2012.00012

Meijer, S., Bramson, B., Toni, I., & Roelofs, K. (2023). Improving approach-avoidance control in social anxiety by targeting phase-amplitude coupling between prefrontal and sensorimotor cortex. Preprint.

Meng, Q., Zhu, Y., Yuan, Y., Ni, R., Yang, L., Liu, J., & Bu, J. (2023). Dual-site beta tACS over rIFG and M1 enhances response inhibition: A parallel multiple control and replication study. Int J Clin Health Psychol, 23(4), 100411. 10.1016/j.ijchp.2023.100411

Mirabella, G., Pani, P., & Ferraina, S. (2011). Neural correlates of cognitive control of reaching movements in the dorsal premotor cortex of rhesus monkeys. J Neurophysiol, 106(3), 1454–1466. 10.1152/jn.00995.2010

Mormann, F., Fell, J., Axmacher, N., Weber, B., Lehnertz, K., Elger, C. E., & Fernandez, G. (2005). Phase/amplitude reset and theta-gamma interaction in the human medial temporal lobe during a continuous word recognition memory task. Hippocampus, 15(7), 890–900. 10.1002/hipo.20117

Nikulin, V. V., & Brismar, T. (2006). Phase synchronization between alpha and beta oscillations in the human electroencephalogram. Neuroscience, 137(2), 647–657. 10.1016/j.neuroscience.2005.10.031

Oldfield, R. C. (1971). The assessment and analysis of handedness: the Edinburgh inventory. Neuropsychologia, 9(1), 97–113. 10.1016/0028-3932(71)90067-4

Oostenveld, R., Fries, P., Maris, E., & Schoffelen, J. M. (2011). FieldTrip: Open source software for advanced analysis of MEG, EEG, and invasive electrophysiological data. Comput Intell Neurosci, 2011, 156869. 10.1155/2011/156869

Pogosyan, A., Gaynor, L. D., Eusebio, A., & Brown, P. (2009). Boosting Cortical Activity at Beta-Band Frequencies Slows Movement in Humans. Current Biology, 19(19), 1637–1641. 10.1016/j.cub.2009.07.074

Reinhart, R. M. G., & Nguyen, J. A. (2019). Working memory revived in older adults by synchronizing rhythmic brain circuits. Nat Neurosci, 22(5), 820–827. 10.1038/s41593-019-0371-x

Schaum, M., Pinzuti, E., Sebastian, A., Lieb, K., Fries, P., Mobascher, A., Jung, P., Wibral, M., & Tüscher, O. (2020). Cortical network mechanisms of response inhibition. Preprint.

Schwab, B., Penen, S. H., & Gann, M. (2025). Connectivity modulation by dual-site tACS – computational and experimental perspectives. Brain Stimulation, 18(1), 249. 10.1016/j.brs.2024.12.107

Seo, J., Lee, J., & Min, B. K. (2024). Out-of-phase transcranial alternating current stimulation modulates the neurodynamics of inhibitory control. NeuroImage, 292, 120612. 10.1016/j.neuroimage.2024.120612

Stinear, C. M., Coxon, J. P., & Byblow, W. D. (2009). Primary motor cortex and movement prevention: where Stop meets Go. Neurosci Biobehav Rev, 33(5), 662–673. 10.1016/j.neubiorev.2008.08.013

Swann, N., Tandon, N., Canolty, R., Ellmore, T. M., McEvoy, L. K., Dreyer, S., DiSano, M., & Aron, A. R. (2009). Intracranial EEG reveals a time- and frequency-specific role for the right inferior frontal gyrus and primary motor cortex in stopping initiated responses. J Neurosci, 29(40), 12675–12685. 10.1523/JNEUROSCI.3359-09.2009

Swann, N. C., Cai, W., Conner, C. R., Pieters, T. A., Claffey, M. P., George, J. S., Aron, A. R., & Tandon, N. (2012). Roles for the pre-supplementary motor area and the right inferior frontal gyrus in stopping action: electrophysiological responses and functional and structural connectivity. NeuroImage, 59(3), 2860–2870. 10.1016/j.neuroimage.2011.09.049

Tan, J., Iyer, K. K., Nitsche, M. A., Puri, R., Hinder, M. R., & Fujiyama, H. (2022). The Effects of Dual-Site Beta tACS over the rIFG and preSMA on Response Inhibition in Young and Older Adults. Preprint.

Thielscher, A., Antunes, A., & Saturnino, G. B. (2015). Field modeling for transcranial magnetic stimulation: A useful tool to understand the physiological effects of TMS? Annu Int Conf IEEE Eng Med Biol Soc, 2015, 222–225. 10.1109/EMBC.2015.7318340

Verbruggen, F., Aron, A. R., Band, G. P., Beste, C., Bissett, P. G., Brockett, A. T., Brown, J. W., Chamberlain, S. R., Chambers, C. D., Colonius, H., Colzato, L. S., Corneil, B. D., Coxon, J. P., Dupuis, A., Eagle, D. M., Garavan, H., Greenhouse, I., Heathcote, A., Huster, R. J., . . . Boehler, C. N. (2019). A consensus guide to capturing the ability to inhibit actions and impulsive behaviors in the stop-signal task. eLife, 8, e46323. 10.7554/eLife.46323

Vinck, M., Oostenveld, R., van Wingerden, M., Battaglia, F., & Pennartz, C. M. (2011). An improved index of phase-synchronization for electrophysiological data in the presence of volume-conduction, noise and sample-size bias. NeuroImage, 55(4), 1548–1565. 10.1016/j.neuroimage.2011.01.055

Wagner, J., Wessel, J. R., Ghahremani, A., & Aron, A. R. (2018). Establishing a Right Frontal Beta Signature for Stopping Action in Scalp EEG: Implications for Testing Inhibitory Control in Other Task Contexts. J Cogn Neurosci, 30(1), 107–118. 10.1162/jocn_a_01183

Weinrich, C. A., Brittain, J. S., Nowak, M., Salimi-Khorshidi, R., Brown, P., & Stagg, C. J. (2017). Modulation of Long-Range Connectivity Patterns via Frequency-Specific Stimulation of Human Cortex. Curr Biol, 27(19), 3061–3068 e3063. 10.1016/j.cub.2017.08.075

Wessel, J. R. (2020). beta-Bursts Reveal the Trial-to-Trial Dynamics of Movement Initiation and Cancellation. J Neurosci, 40(2), 411–423. 10.1523/JNEUROSCI.1887-19.2019

Wessel, J. R., & Aron, A. R. (2013). Unexpected events induce motor slowing via a brain mechanism for action-stopping with global suppressive effects. J Neurosci, 33(47), 18481–18491. 10.1523/JNEUROSCI.3456-13.2013

Wessel, J. R., Jenkinson, N., Brittain, J. S., Voets, S. H., Aziz, T. Z., & Aron, A. R. (2016). Surprise disrupts cognition via a fronto-basal ganglia suppressive mechanism. Nat Commun, 7, 11195. 10.1038/ncomms11195

Wessel, J. R., Waller, D. A., & Greenlee, J. D. (2019). Non-selective inhibition of inappropriate motor-tendencies during response-conflict by a fronto-subthalamic mechanism. eLife, 8, e42959. 10.7554/eLife.42959

Woods, A. J., Antal, A., Bikson, M., Boggio, P. S., Brunoni, A. R., Celnik, P., Cohen, L. G., Fregni, F., Herrmann, C. S., Kappenman, E. S., Knotkova, H., Liebetanz, D., Miniussi, C., Miranda, P. C., Paulus, W., Priori, A., Reato, D., Stagg, C., Wenderoth, N., & Nitsche, M. A. (2016). A technical guide to tDCS, and related non-invasive brain stimulation tools. Clin Neurophysiol, 127(2), 1031–1048. 10.1016/j.clinph.2015.11.012

