## Supplementary_materials for "In-phase and anti-phase dual-site beta tACS differentially influence functional connectivity and motor inhibition": Supplemantary material.docx

*Corresponding author

### VAS report

Participants reported significantly higher tiredness levels after the experiment compared to before (Z = -4.722, p < 0.001; Table S1). No major adverse effects or stimulation-related complications were observed. All participants perceived the stimulation as being applied bilaterally. Two participants mentioned that the stimulation did not feel continuous. Regarding discomfort, only one participant reported strong itching. Others experienced no or mild levels of itching or warming sensations. Additionally, two participants reported a metallic or iron taste during stimulation.

Table S1. VAS scores

| **VAS items** | Rating (10-point) | Wilcoxon Signed-Rank Test | **Sensation^#^** | None | Mild | Moderate | Severe |
| --- | --- | --- | --- | --- | --- | --- | --- |
| tiredness-pre | 2.741 ± 1.868 | Z = 4.050, p < 0.001***** | itching | 16 | 18 | 4 | 1 |
| tiredness-post | 4.738 ± 2.136 |  | burning | 30 | 8 | 1 | 0 |
| intensity | 5.603 ± 1.603 | Z = -1.563, p = 0.118 | warmth | 17 | 17 | 5 | 0 |
| focality | 5.179 ± 1.823 | Z = 0.593, p = 0.553 | Metallic/iron taste | 36 | 2 | 1 | 0 |
| pain | 2.692 ± 1.580 | Z = -1.106, p = 0.269 | fatigue/decreased alertness | 8 | 13 | 17 | 1 |
| comfort | 4.436 ± 1.836 | Z = 1.376, p = 0.169 |  |  |  |  |  |

VAS item scores are presented as mean ± standard deviation. *****p < 0.05. **^#^** The number of participants who reported experiencing the corresponding sensation levels.
